# High-resolution structure of a mercury cross-linked ZIP metal transporter reveals delicate motions and metal relay for regulated zinc transport

**DOI:** 10.1101/2023.04.20.537755

**Authors:** Tuo Zhang, Yao Zhang, Dexin Sui, Jian Hu

## Abstract

Zrt-/Irt-like protein (ZIP) divalent metal transporters play a central role in maintaining trace element homeostasis. The prototypical ZIP from *Bordetella bronchiseptica* (BbZIP) is an elevator-type transporter, but the dynamic motions and detailed transport mechanism remain to be elucidated. Here, we report a high-resolution crystal structure of a mercury-crosslinked BbZIP variant at 1.95 Å, revealing an upward rotation of the transport domain in the new inward-facing conformation and a water-filled metal release channel that is divided into two parallel pathways by the previously disordered cytoplasmic loop. Mutagenesis and transport assays indicated that the newly identified high-affinity metal binding site in the primary pathway acts as a “metal sink” to reduce the transport rate. The discovery of a hinge motion around an extracellular axis allowed us to propose a sequential hinge-elevator-hinge movement of the transport domain to achieve alternating access. These findings provide key insights into the transport mechanisms and activity regulation.

## Introduction

Multiple *d*-block metal elements are essential for life. Among the transporters that control the fluxes of these micronutrients across biological membranes, members of the Zrt-/Irt-like protein (ZIP) family play a central role (*1-3*). The ZIP family comprises divalent *d*-block metal transporters that are ubiquitously expressed in all kingdoms of life. This ancient and highly diverse family mediates metal influx into the cytoplasm either from the extracellular milieu or from intracellular organelles/vesicles. In humans, the fourteen ZIPs (ZIP1-14) are largely involved in zinc (Zn), iron (Fe) and manganese (Mn) homeostasis. Some ZIPs are associated with human diseases (*4-9*) and are therefore considered as potential targets for novel therapies (*10-18*). For example, a monoclonal antibody against human ZIP6 (LIV-1) is in clinical trials for breast cancer and other solid tumors (*19, 20*). ZIP4, which was found to be aberrantly upregulated in many cancers, has been shown to be associated with cancer cell growth and metastasis (*21-29*).

Recent structural studies have provided important insights into the transport mechanism of the ZIPs. The structure of a prototypic ZIP from *Bordetella bronchiseptica* (BbZIP) revealed the structural framework of the transmembrane domain that is conserved in the entire ZIP family (*30*). An elevator-type transport mode for BbZIP was later proposed based on the two-domain architecture, evolutionary covariance analysis, and an outward-facing conformation (OFC) model validated by chemical crosslinking and cysteine accessibility assay (*31, 32*). According to the proposed mechanism, the four transmembrane helix (TM) bundle composed of TM1/4/5/6 (the transport domain), which carries the metal substrate, moves vertically by 8 Å relative to the static and wall-like scaffold domain (TM2/3/7/8) to achieve alternating access. Although several structures of the inward-facing conformation (IFC) have been solved experimentally and an OFC model has been proposed, the upward movement of the transport domain relative to the scaffold domain, which is an essential step for the proposed conformational transition from the IFC to the OFC, has not been observed. Rather, by comparing the transporter in the apo (metal-free) state with the metal-bound state, two independent studies have revealed a downward movement of the transport domain, which is associated with metal release from the transport site (*31, 32*).

In this work, we report a 1.95 Å crystal structure of a BbZIP double cysteine variant in a new inward-facing conformation stabilized by mercury (Hg) cross-linking, revealing an upward rotation of the transport domain, which, together with previously solved BbZIP structures, indicates a continuous hinge motion. The high-resolution structure also reveals a new high-affinity metal site involving the previously disordered cytoplasmic histidine-rich loop in the water-filled metal release pathway. Mutagenesis and functional studies on BbZIP and human ZIP4 (hZIP4) indicate that the newly identified metal binding site acts as a “metal sink” to limit metal release to the cytoplasm. We believe that these new findings provide key insights into the metal transport mechanism of the ZIP family, which may shed light on the mechanistic studies of other transition metal transporters.

## Results and Discussion

### Crystallization of a BbZIP variant cross-linked by Hg^2+^ in the absence of free Cd^2+^

In an effort of locking the transporter in an OFC of BbZIP, we identified a double cysteine variant (A95C/A214C) that can be cross-linked by HgCl_2_ (**Figure 1**). In the previously solved BbZIP structures, the distances between the C_β_ atoms of the two residues vary in the range of 7.4-11.8 Å, which are longer than the minimum distance required for a Hg-mediated cross-linking (7.0 Å). We expected that the cross-linked variant would adopt a conformation different from any previously solved structures. Since the addition of Hg^2+^ to the purified proteins caused a clear gel shift in non-reducing SDS-PAGE only for the double variant but not for any of the single variants (A95C and A214C), the gel shift was completely abolished by treatment with 2-mercaptoethanol, and pretreatment with a thiol reacting agent N-ethylmaleimide (NEM) prevented cross-linking, we concluded that the two introduced cysteine residues, which are the only cysteine residues in the variant due to the lack of endogenous cysteine in the wild-type BbZIP, must have been cross-linked by Hg^2+^ via two Hg-S bonds. We found that the Hg-mediated cross-linking stabilized the protein in solution even when free cadmium (Cd^2+^) was removed by a quick treatment with EDTA followed immediately by desalting. We then crystallized the cross-linked variant in lipidic cubic phase (LCP) under a condition with no Cd^2+^ included in the crystallization buffer. As the previous BbZIP crystals were obtained either in the presence of high concentration of Cd^2+^ (tens to hundreds of millimolar) or at low pH (pH 3-4), this new crystal form allowed us to reveal a new IFC and revisit metal binding to BbZIP.

**Figure 1.**
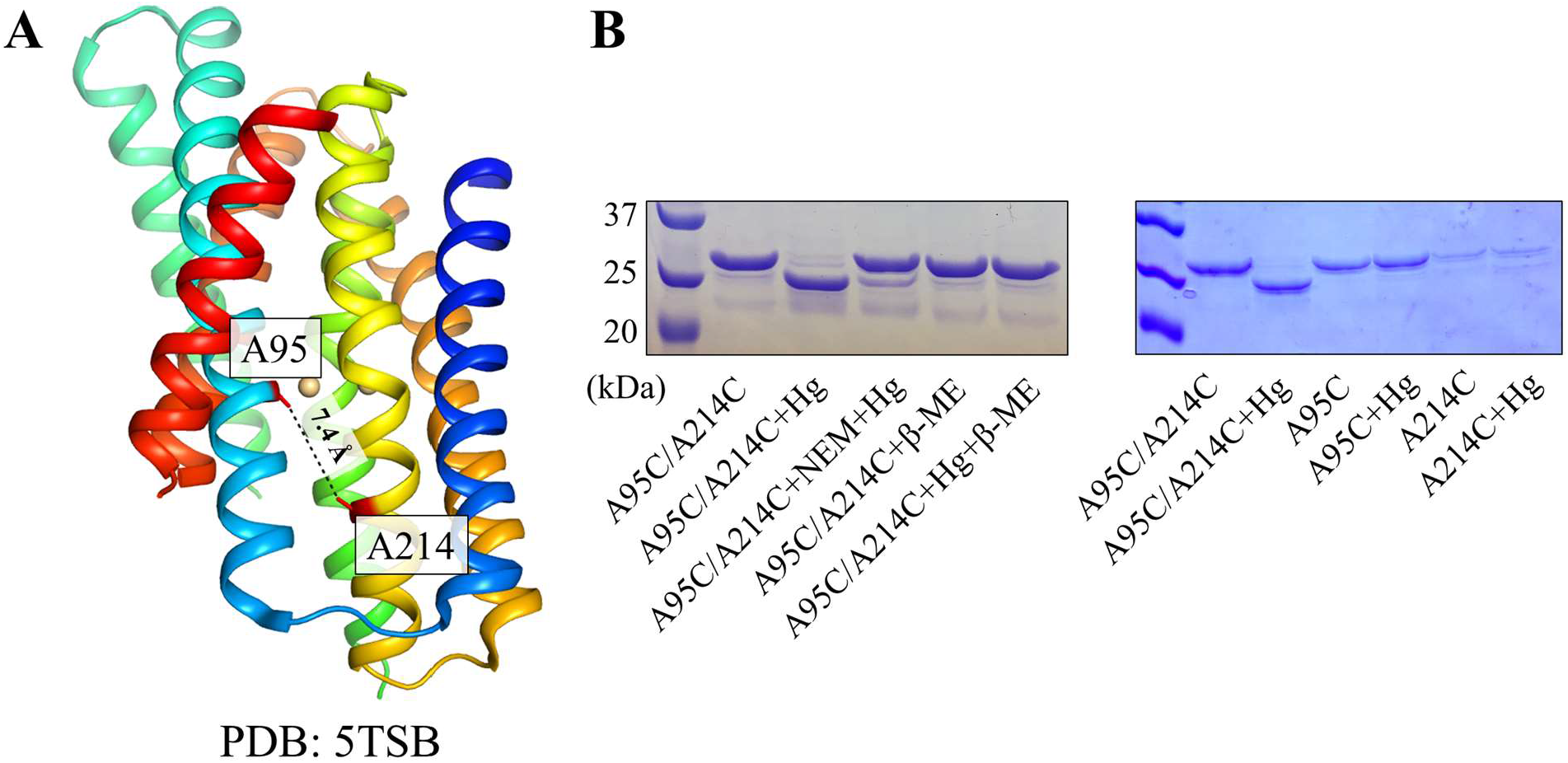
Hg^2+^-mediated cross-linking of the A95C/A214C variant. (**A**) A95 and A214 in IFC of BbZIP (PDB: 5TSB). The distance between the C_β_ atoms (7.4 Å) is greater than the minimum distance required for an S-Hg-S linkage (7.0 Å). (**B**) Cross-linking of the variants of A95C/A214C, A95C, and A214C by HgCl_2_ at the 1:10 molar ratio. Pretreatment with N-ethylmaleimide (NEM) or post treatment with 2-mercaptoethanol (β-ME) prevents or reverses the cross-linking reaction.

### Overall structure of the Hg^2+^ cross-linked A95C/A214C variant

The crystal structure was solved at 1.95 Å, representing the highest resolution for BbZIP (**Figure 2A, Table S1**). As revealed in the structures of BbZIP in the apo state (*31, 32*), the transporter consists of nine TMs with the N-terminal TM0 only weakly associated with TM3 and TM6. The variable interface between TM0 and the rest of the transporter strongly indicates that TM0 is unlikely to play a crucial role in transporter function. Different from our previous structure solved in the apo state (PDB: 8CZJ), the very N-terminal amphipathic helix α0a was not resolved in the new structure, highlighting the structural flexibility of this highly variable region. The conserved eight TMs is composed of two domains – TM1/4/5/6 form the transport domain and TM2/3/7/8 form the scaffold domain, according to the proposed elevator-type transport mode (*31, 32*). As expected, a Hg^2+^ was found to coordinate with the two introduced cysteine residues (C95 from TM2 and C214 from TM5) with an ideal geometry for a S-Hg-S linkage (**Figure 3**), fixing the relative orientation between the transport domain and the scaffold domain, which likely accounts for the high resolution of this crystal structure. As the distance between the two C_β_ atoms has been shorten to 6.9 Å, the protein adopts an IFC different from those seen in the previously solved structures, which will be discussed in a later section.

**Figure 2.**
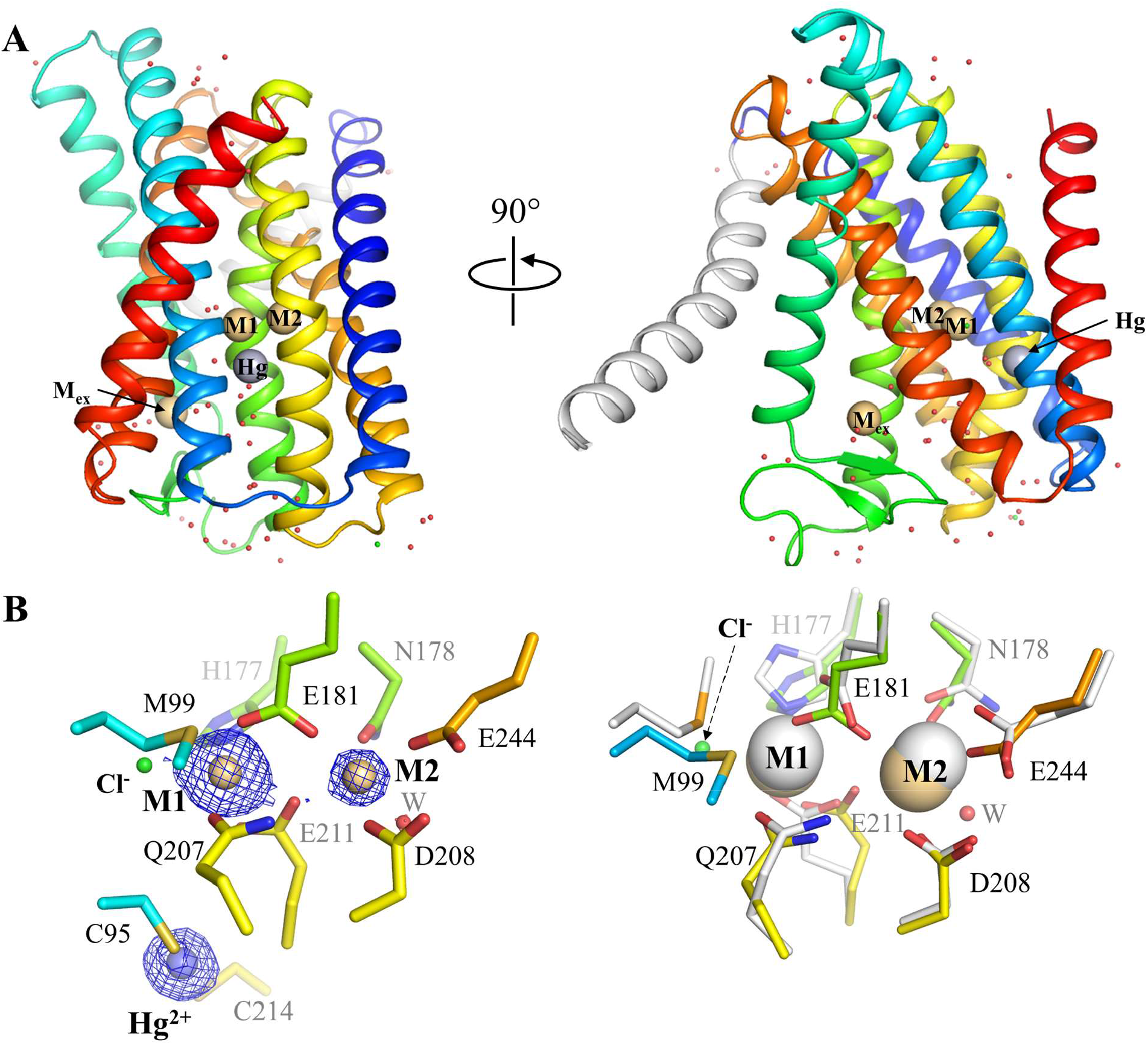
Structure of the Hg^2+^ cross-linked A95C/A214C variant. (**A**) Overall structure. The conserved eight TMs are colored in rainbow and TM0 is in grey. Cd^2+^ at the M1, M2, and M_ex_ sites are depicted as light-brown spheres. Hg^2+^ is shown as a grey sphere. (**B**) The structure of the BMC. *Left*: the BMC and Hg^2+^ in the new structure. The blue meshes indicate the electron densities of Cd^2+^ and Hg^2+^ (2Fo-Fc map, σ=4). The occupancy of Cd^2+^ at the M2 site (0.42) is half of that at the M1 site (0.85). *Right*: structural comparison of the BMC in the new structure (rainbow color) with the Cd^2+^ bound structure (white, PDB: 5TSB). Note that a chloride ion (green sphere) is in the position which is occupied by the sulfur atom of M99 in 5TSB.

**Figure 3.**
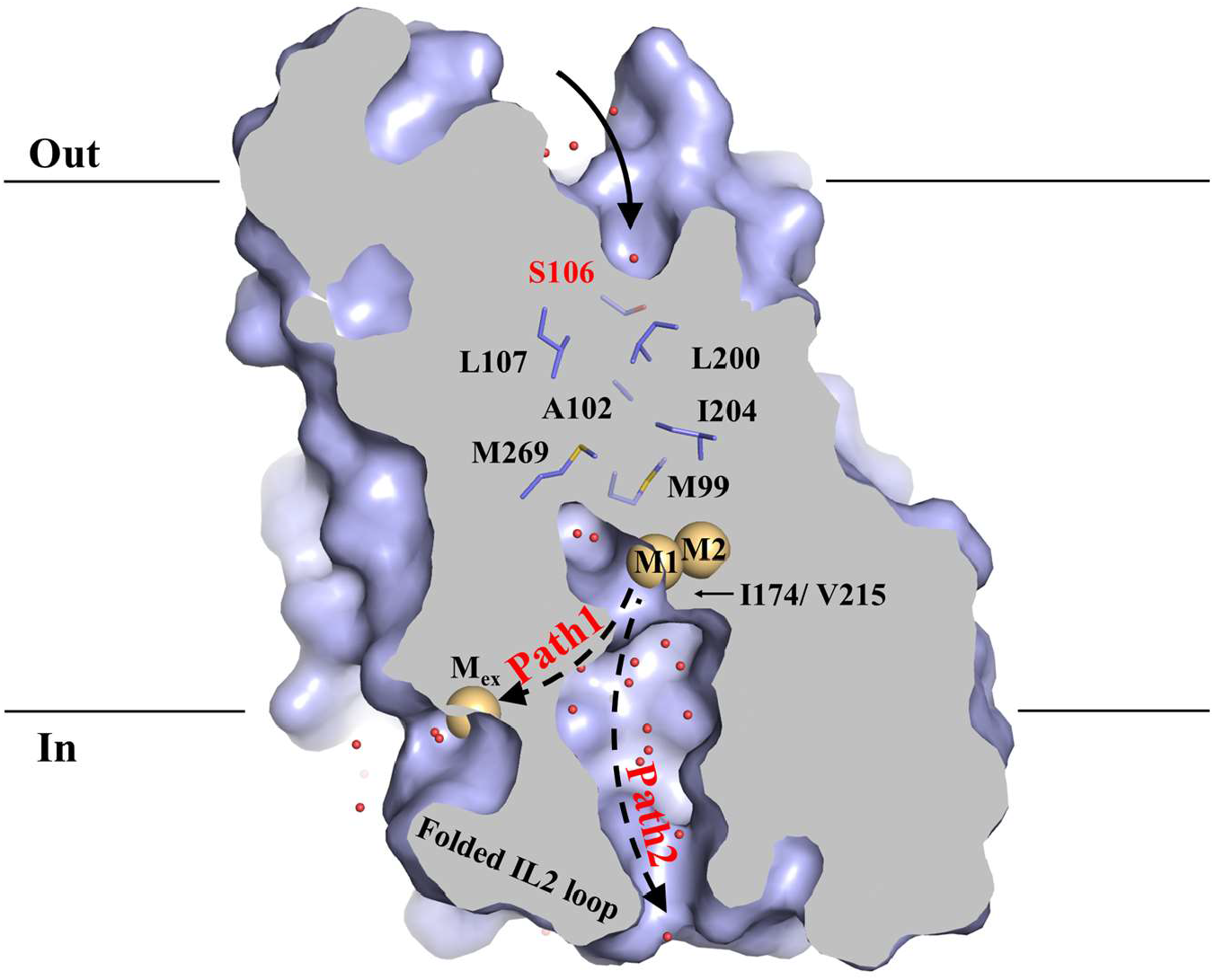
Water molecules along the transport pathway. Cross-section view of the transporter in surface mode reveals a blocked pathway toward extracellular space (by the indicated non-polar residues) and a water-filled pathway toward the cytoplasm. S106 (red) is located at the entrance of the transport pathway (indicated by the curved solid black arrow). Water molecules are depicted as red spheres. Three bound Cd^2+^ (at M1, M2, and M_ex_) are shown as light-brown spheres. The dashed arrows indicate two potential pathways (Path1 and Path2), which are separated by the folded IL2 loop, for metal release from the M1 site to the cytoplasm.

### The binuclear metal center (BMC) at the transport site

Although the cross-linked variant was crystallized in the absence of free Cd^2+^, the two closely associated metal binding sites (M1 and M2) are still occupied by Cd^2+^ ions (**Figure 2B**), indicating that metal binding to the BMC is not an artifact. Occupancy refinement of the two bound metal ions showed that the occupancy of M1 (0.85) is twice of that of M2 (0.42), indicating that M1 has a higher affinity than that of M2. This result is consistent with the previous report that the occupancy of metal at the M2 site was decreased significantly whereas that at the M1 site did not when the Cd^2+^-bound crystals were soaked in a metal-free solution (*33*).

As the metal binding sites are exclusively located in the transport domain, for better comparing the coordination environment of the bound metals, the transport domain of the new structure was aligned with that of the previously solved Cd-bound structure (PDB: 5TSB, **Figure 2B**). Although the overall arrangement of the BMC is similar to that shown in previous structures, some differences were still evident. Particularly, a chloride ion was found to coordinate to Cd^2+^ at the M1 site. A highly ordered water molecule was excluded because of the otherwise residual positive density in the Fo-Fc map (**Figure S1)**. After examining possible ligands present in the crystallization buffer, only a chloride ion would match the density with a *B* factor comparable to that of the metal at M1. As there is no evidence supporting metal co-transport with chloride ion and there is no residue coordinating to the chloride ion, it is unlikely that chloride ion is a substrate of BbZIP. Rather, the negative charge of chloride ion may assist metal release from the high-affinity transport site. This may be part of the mechanism that allows efficient metal release from the high-affinity transport site. As the chloride ion occupies the position which was occupied by a coordinating sulfur atom from the side chain of M99, the latter swings away from the M1 metal and adopts a rotamer conformation similar to that observed in the previously solved zinc-bound structure (PDB: 5TSA). Likely due to the negative charge of the chloride ion, the side chain of E211 is displaced to a position between the two metals, forming another bridging residue in addition to E181. Overall, the coordination sphere of the M1 metal underwent a greater change than that of the M2 metal, and the structural changes of the residues involved in metal chelation revealed conformational plasticity of the transport site.

### Water molecules along the transport pathway

The high-resolution structure allowed us to visualize the ordered water molecules around the protein, and particularly, along the transport pathway (**Figure 3**). As the transporter is in an IFC, the metal release pathway is open and filled with water molecules. A couple of waters are found to penetrate the transport site to reach the bottom of the sealed pathway toward the extracellular space. Accordingly, hydrophobic residues including M99 and M269 are water accessible, which is consistent with the results of a hydroxyl radical footprinting experiment (*34*). There is a clear water-filled pathway for the M1 metal to release to the cytoplasm, whereas the pathway for the M2 metal to reach the cytoplasm is blocked by I174 and V215. This explains why the Cd^2+^ bound at the M1 site was ready to be replaced by a Zn^2+^ when the Cd-bound crystals were soaked with a Zn^2+^-containing solution whereas the Cd^2+^ at the M2 site could not (*30*). Consistent with the IFC, waters from the extracellular side can only reach S106 at the entrance of the transport pathway, leaving an 8-9 Å pathway blocked by hydrophobic residues (**Figure 3**).

### A new metal binding site at the exit of the transport pathway

Different from other IFC structures, the metal release channel in the new structure is divided into two parallel pathways by the folded cytoplasmic loop connecting TM3 and TM4 (intracellular loop 2, IL2) – one pathway is mediated by a new metal binding site (M_ex_) located at the lipid-water interface (Path1), whereas the other allows the M1 metal to be released directly to the cytoplasm (Path2) (**Figure 3**). The metal at the M_ex_ site is coordinated with two histidine residues (H149 and H151) from the IL2, D144 from TM3, E276 from TM7, and a water molecule, forming a distorted octahedral coordination sphere (**Figure 4**). The occupancy of the metal at the M_ex_ site (0.86) is nearly identical to that of the M1 metal, indicating that it is a high-affinity metal binding site. However, this metal binding site was not observed in the previous structures of BbZIP, which can be attributed to the dynamic nature of the IL2. In the new structure, the IL2 folds into two short anti-parallel β-strands and the entire loop is further stabilized by hydrogen bonds involving the residues of R166, T173, and E280. Similar to the counterpart of the IL2 in human ZIP4 (*35, 36*), the IL2 is the longest cytoplasmic loop in BbZIP and predicted to be intrinsically disordered (**Figure S2**). As folding of intrinsically disordered region is sensitive to local environment, it is plausible that the IL2 loop will fold differently or become unfolded when metal substrate is released to the cytoplasm and/or the transporter adopts different conformations.

**Figure 4.**
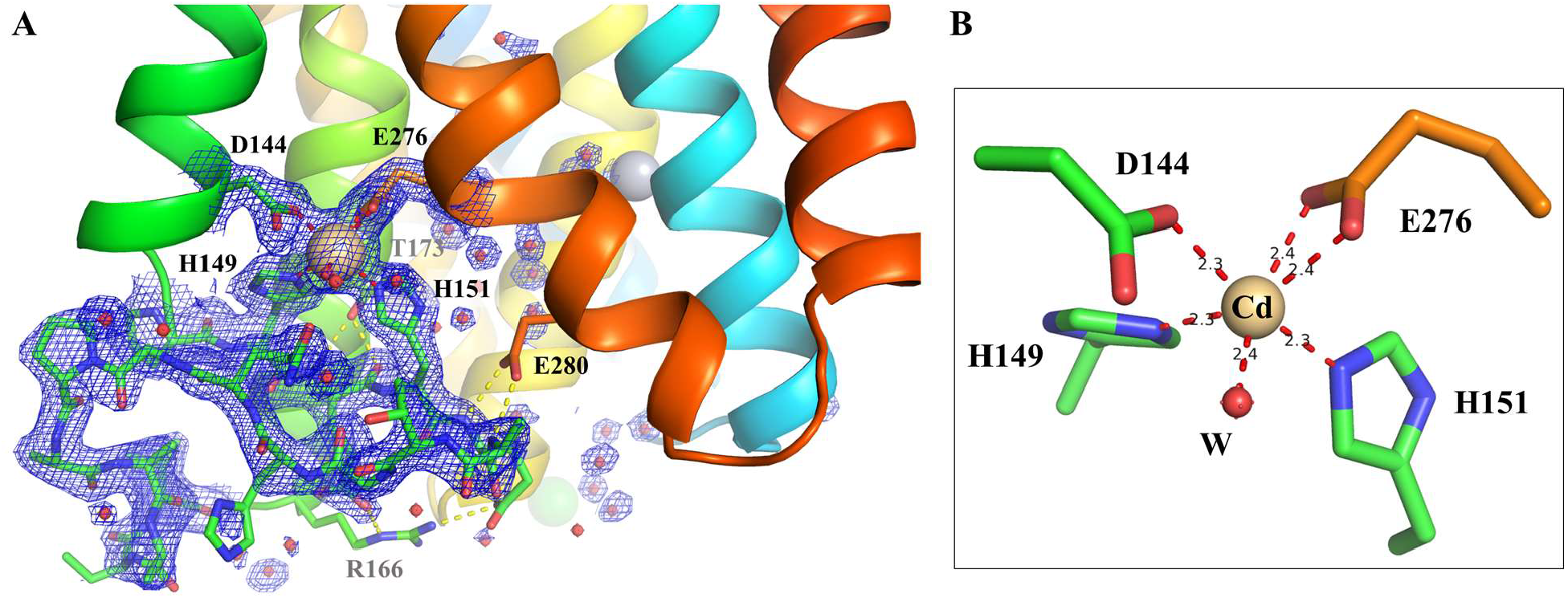
The structure of the IL2 loop and the M_ex_ site at the exit of the transport pathway. (**A**) 2Fo-Fc map (σ=1) of the IL2 loop and water molecules. The residues in the IL2 loop are shown in stick mode and the rest of the protein is in cartoon mode. Water molecules are depicted as red spheres. The hydrogen bonds involved in stabilizing the IL2 loop structure are shown as yellow dashed lines. (**B**) The distorted octahedral coordination sphere of Cd^2+^ at the M_ex_ site. The distances between Cd^2+^ and the ligands are labeled in angstrom.

### Functional assessment of the M_ex_ site

To assess the function of the M_ex_ site, the four residues forming the M_ex_ site were substituted with alanine in three variants – D144A, E276A, and H151A/H154A. The function of BbZIP was evaluated by using an approach adapted from a recent report (*31*). In brief, the wild-type and the variants of BbZIP were expressed in the *E*.*coli* strain C43 (DE3), and the cells were applied to the LB agar plates containing 1.5 mM ZnCl_2_. A high zinc concentration was found to be necessary to see a growth difference between the cells transformed with an empty vector and those expressing BbZIP (**Figure S3**), and the functions of the variants relative to the wild-type protein can be evaluated accordingly. As shown in **Figure 5A**, the cells expressing any of the three variants exhibited suppressed growth similar to those expressing the wild-type BbZIP, suggesting that none of the residues forming the M_ex_ site is essential for BbZIP function. Instead, since the variants, in particular the H149A/H151A variant, achieved the same level of growth suppression as the wild-type protein even though their expression levels are significantly lower than that of the latter as indicated in the Western blot experiment, the zinc transport activities of these variants may have been increased. Accordingly, the role of the M_ex_ site appears to be to retain the metal with the transporter for a longer time before the metal is released into the cytoplasm.

**Figure 5.**
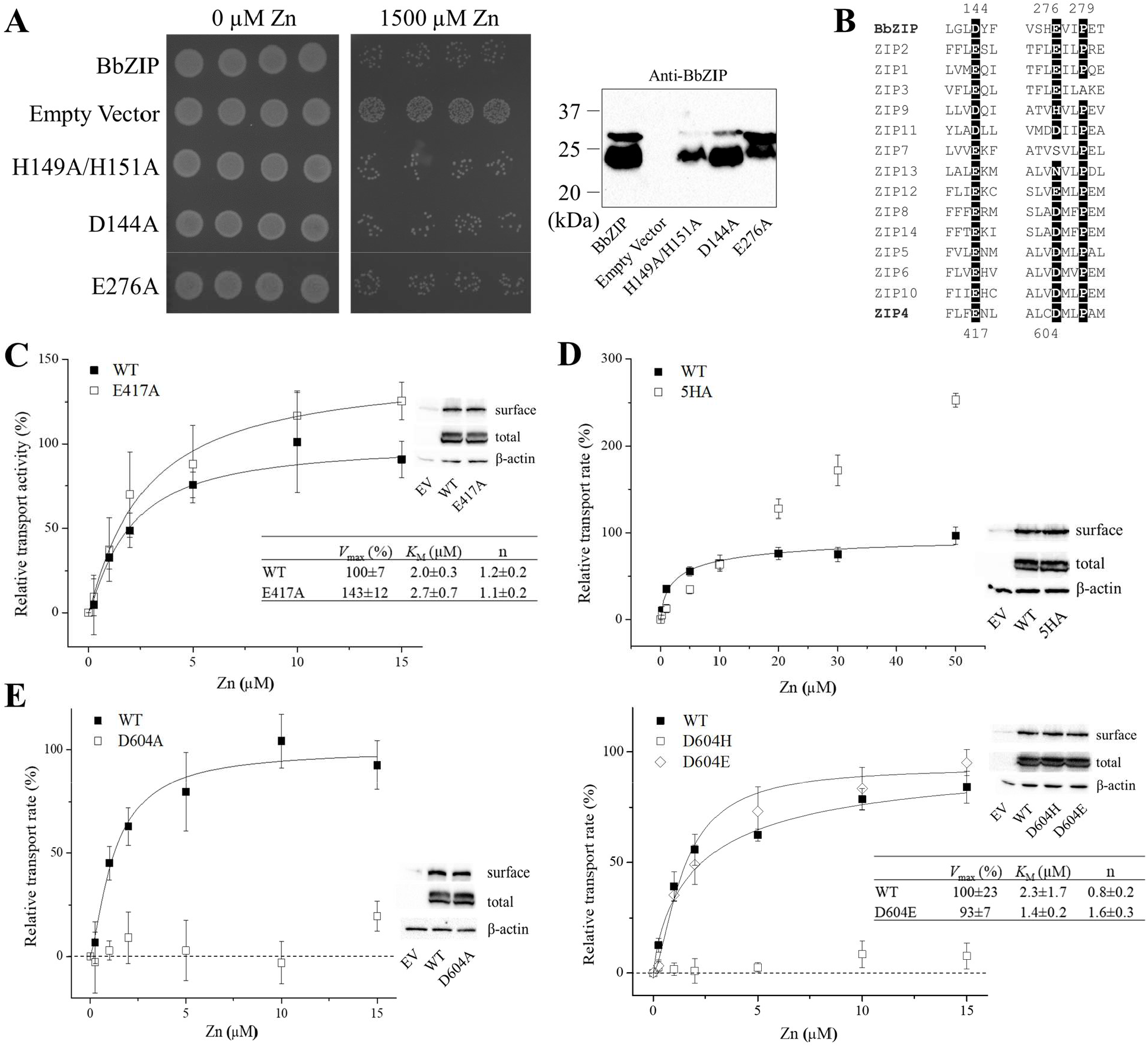
Mutagenesis and functional characterization of the M_ex_ site. (**A**) Growth phenotype-based transport function assessment of BbZIP and the variants. The *E*.*coli* cells (C43 strain) with or without expressing BbZIP or the variants were adjusted to OD_600_=0.001 and 5 μl of the cell suspension for each construct was applied to the LB agar plates. The cells were allowed to grow at 37°C for 24 hours before images were taken. Four replicates were included for each construct. A custom monoclonal antibody was used as the primary antibody to detect BbZIP in Western blot. (**B**) Sequence alignment of BbZIP and human ZIPs. D144, E276, and P279 in BbZIP and their counterparts in other ZIPs are highlighted. (**C**-**E**) Functional study of the hZIP4 variants in the cell-based ^65^Zn transport assay. HEK293T cells transiently transfected with vectors harboring the wild-type or mutated hZIP4 genes were incubated with indicated concentrations of ZnCl_2_ composed of radioactive ^65^Zn and non-radioactive zinc at 37°C for 30 minutes. Transport was terminated by addition of an ice-cold EDTA-containing buffer and the cells were washed extensively before the radioactivity associated with cells was quantified by using a gamma counter. Zinc transport activity was calculated by subtracting the radioactivity associated with the cells transfected with the empty vector and further calibrated by hZIP4 cell surface expression level determined in Western blot experiments. The relative transport activity of a variant was calculated by comparing with that of the wild-type hZIP4. The dose curves were fitted with the Hill model and the obtained kinetic parameters are listed in the tables. Data shown are representative of one of three independent experiments. Three replicates were included for each data point in one experiment. The error bars indicate S.D.

Next, taking advantage of the fact that the transport kinetic parameters of human ZIP4 (hZIP4) can be determined by using an established transport assay (*37-41*), we tested the functions of the residues in hZIP4 that are topologically equivalent to those forming the M_ex_ site in BbZIP (**Figure 5B**). E417, which is equivalent to D144 in BbZIP, was found to be dispensable for activity (**Figure 5C**). Instead, the E417A variant exhibited a modestly increased *V*_max_ when compared to the wild-type hZIP4. Different from the short IL2 of BbZIP, the corresponding loop of hZIP4 is longer and harbors five histidine residues. We therefore replaced all the histidine residues with alanine to generate the 5HA variant. Unexpectedly, the 5HA variant exhibited a drastically changed kinetic properties (**Figure 5D**). The dose-dependent profile appeared to be linear at low zinc concentrations and curved only at high zinc concentrations. Because the curve did not saturate in the range of zinc concentrations tested in this work, we could not fit the curve to obtain the value of *K*_M_ or *V*_max_. However, the curve profile suggested that both *K*_M_ and *V*_max_ increased significantly. As the 5HA variant showed no difference in the overall expression or cell surface expression when compared to the wild-type protein, the increased *V*_max_ indicated that the turnover rate of the transporter was increased significantly. The greatly increased transport activity upon histidine substitution by alanine in the IL2 is reminiscent of AtMTP1, a vacuolar Zn^2+^/H^+^ antiporter from *Arabidopsis thaliana* belonging to the cation diffusion facilitator (CDF) superfamily (*42*), whose activity is greatly stimulated when the cytoplasmic his-rich loop was deleted. Similarly, replacing the very C-terminal cytoplasmic “His-Cys-His” metal binding motif of the copper transporter CTR1 with alanine residues greatly increased both *K*_M_ and turnover rate (*43*), a behavior analogous to the 5HA variant of hZIP4. Although it is unclear whether the metal binding motifs in these metal transporters are able to form a M_ex_-like high-affinity binding site within the metal translocation pathway, the use of a “metal sink” appears to be a frequently used strategy to regulate transport activities of *d*-block metal transporters. Different from the functionally dispensable E276 in BbZIP, the corresponding residue in hZIP4, D604, was found to be essential for the zinc transport activity of hZIP4 (**Figure 5E**), Substitution of D604 with alanine completely abolished the zinc transport activity, and only the D604E variant, not the D604H variant, showed activity similar to that of wild-type hZIP4.

Collectively, eliminating the M_ex_ site in BbZIP or the putative equivalent in hZIP4 did not diminish transporter’s function. Rather, the M_ex_ site (and its putative equivalent in hZIP4) reduces zinc transport by acting as a “metal sink” to limit metal release into the cytoplasm. Although E276 is functionally dispensable in BbZIP, the equivalent residue D604 in hZIP4 is essential for activity, suggestive of additional roles of D604 in hZIP4, which is discussed in the next section.

### Metal relay from the BMC to the cytoplasm

To understand the structural basis of the different functions of D144 and E276, metal binding to these residues were surveyed in the available metal-bound structures (**Figure S4**). D144 was consistently observed to participate in metal binding in the structures of 5TSB, 5TSA, 7Z6M, and the new structure reported in this work; in contrast, in addition to participating in the M_ex_ site as shown in this work, E276 was only found to bind a low-occupancy zinc ion when the Cd-bound crystals were soaked with a high concentration of zinc ions (100 mM) (*33*). To better understand the function of E276, three structures (5TSA, 7Z6B, and the new structure) are superimposed. As shown in **Figure 6**, a clear pathway from the BMC to the cytoplasm is visualized, through which the metal substrate likely translocates in a stepwise manner. Since the M2 site has no direct access to the cytoplasm (blocked by I174 and V215, **Figure 3**), metal release from the BMC must start from the M1 site. Swinging of the side chain of H177 away from the M1 site opens a pathway for metal to leave the high-affinity transport site. The metal is then handed to E276, which forms a transient metal binding site with H177 as shown in 5TSA. Next, as revealed in the superimposed structures, E276 undergoes a nearly right-angle rotation, directing the metal substrate to the M_ex_ site (“metal sink”, Path1) where it is retained before being slowly released into the cytoplasm. As such, E276 appears to play multiple roles – it transiently holds the metal that leaves the M1 site; its structural flexibility allows it to guide the metal to enter the M_ex_ site; and it joins the M_ex_ site to keep the metal with the transporter. The conserved P279 as a helix breaker and the intrinsic high flexibility of the segment connecting TM7 and TM8 (**Figure S2**) likely account for the significant dynamics of E276. As Path2 lacks metal-chelating residues and is rich in hydrophobic residues (**Figure S5**), Path1 may represent the main metal release pathway for the wild-type transporter. Consistently, when Path1 (or the equivalent in hZIP4) is eliminated by mutations, the transport rate is increased because the metal ion held at the transient binding site directly enters and diffuses through the water-filled Path2 (**Figures 3**&6). The putative function of the M_ex_ site is a kinetic trap, meaning that the energy barrier into it is low but out of it is high, which would explain why Path1, rather than Path2 which allows for a greater overall transport rate, is the main metal release pathway.

**Figure 6.**
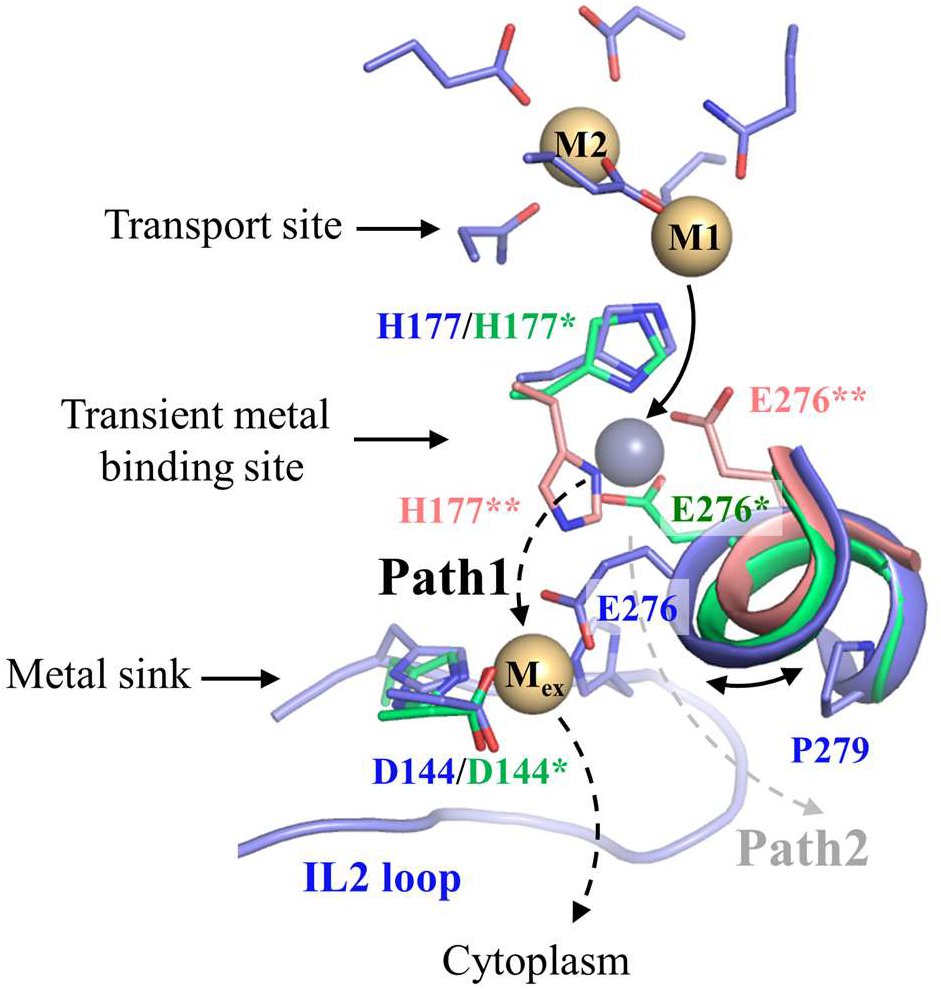
Metal relay in the metal release pathway. The new structure (blue), the Cd-bound structure (PDB: 7Z6M, green), and the zinc-substituted structure (PDB: 5TSA, pink) are superimposed to reveal the metal relay. Cd^2+^ and Zn^2+^ are shown in light brown spheres, respectively. Cd^2+^ ions in the new structure and 7Z6M are generally superimposable and the transient binding site occupied by a Zn^2+^ is only visible in 5TSA. In Path1 (the primary pathway, indicated by the black dashed arrows), a nearly 90-degree rotation of E276, which is enabled because of the flexibility of the cytoplasmic side of TM7 (shown as a segment of an α-helix), directs metal substrate from the transient binding site to the M_ex_ site (metal sink) before metal is released into the cytoplasm. When the M_ex_ is eliminated by mutations, metal released from the transient binding site may enter the cytoplasm via the alternative Path2 (the dashed arrow in grey).

Although the E276A mutation did not reduce BbZIP function, D604 in hZIP4 is indispensable for zinc transport activity. We postulate that metal release from the BMC of BbZIP is not absolutely dependent on E276, whereas it is crucial for hZIP4 to use D604 and probably additional metal chelating residues to prevent the metal substrate from being pulled back to the high-affinity transport site. The lack of transport activity of the D604H variant highlights the importance of the side chain’s flexibility and implies a restricted geometry of the putative transient metal binding site in hZIP4. Consistent with the proposed function of D604 in metal binding, the D604-equivalent residue in human ZIP2, E276, was shown to be essential for Cd^2+^ transport (*44*). Replacing E276 in ZIP2 with alanine nearly completely abolished activity, whereas the transport active E276Q variant showed a 3-fold increase in *K*_M_. The results from human ZIPs suggest a conserved mechanism even though ZIP4 and ZIP2 are members of the LIV-1 and the ZIPII subfamilies, respectively.

### Delicate motions of the transport domain

When compared with the structures of BbZIP in the metal bound state, the metal-free state structures consistently showed a downward rotation of the transport domain relative to the scaffold domain (*31, 32*), whereas the new Hg^2+^ cross-linked structure revealed an upward rigid-body rotation of the transport domain relative to the scaffold domain (**Figure 7A**). Notably, when the scaffold domains of the representative IFC structures are aligned, a continuous displacement of the transport domain can be visualized – it undergoes a hinge motion around a fixed axis that is located at the extracellular terminus of TM6 (referred to as the extracellular axis). One might expect that extrapolation of this rotation would allow the transporter to switch from the IFC to the OFC. However, since further rotation of the transport domain would inevitably force the polar/charged residues on the cytoplasmic side into the lipid bilayer, this rotation alone cannot complete the IFC-to-OFC transition. To profile the conformational changes in a transport cycle, we applied the approach of repeat-swap homology modeling, which has been used to produce an experimentally validated OFC model of BbZIP in our previous study (*32*), to generate additional OFCs by using the structure reported in this work and the metal-free state structure (8CZJ) as templates (**Figure S6**). Structural comparison of the generated OFC models showed that the transport domain undergoes a hinge motion around the axis that is located at the cytoplasmic end of TM1 (referred to as the cytoplasmic axis), which mirrors the hinge motion of the transport domain in IFCs. Taken together, a trajectory of the sequential movements of the transport domain is proposed and illustrated in **Figure 7B**. Metal substrate(s) enter the BMC via the open pathway in the most OFC, after which the transport domain rotates clockwise around the cytoplasmic axis to narrow and eventually close the pathway (occluded state). Then, the transport domain undergoes a canonical elevator motion by sliding vertically down against the scaffold domain to switch to an IFC. Lastly, a counterclockwise hinge motion of the transport domain around the extracellular axis occurs, allowing metal release from the disassembled M1 site, completing the OFC-to-IFC transition and metal transport from the extracellular space to the cytoplasm. Upon comparison with known elevator-type transporters (*45*), we found that BbZIP is unique in that the elevator motion of the transport domain is preceded and followed by two mirrored hinge motions that are likely functionally coupled to the binding/release of metal substrates.

**Figure 7.**
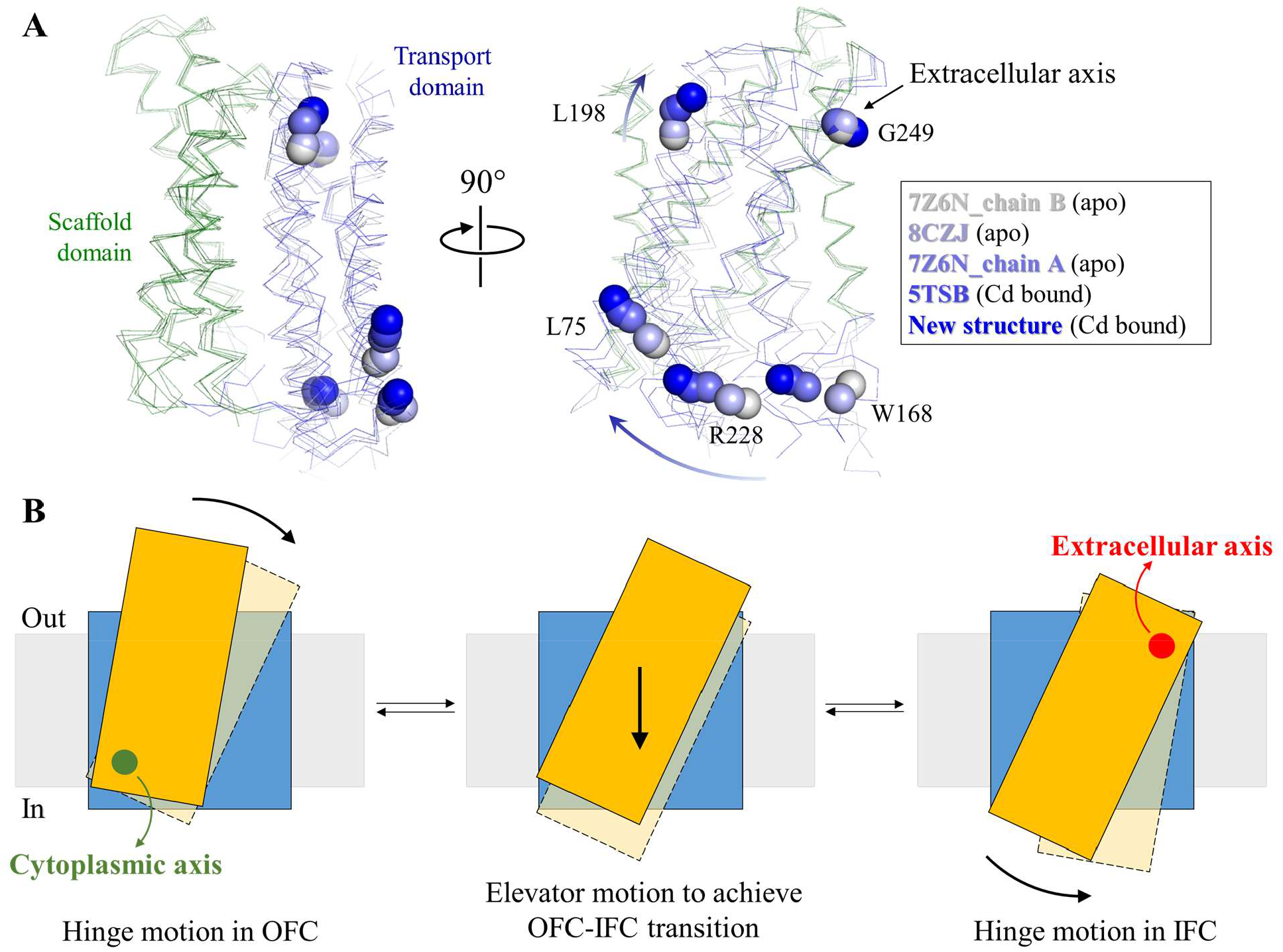
Proposed conformational changes of the transport domain during transport. (**A**) Hinge motion of the transport domain around the extracellular axis revealed in the superimposed IFC structures. The scaffold domains (green) are structurally aligned. The transport domain colored from light blue to dark blue are shown in ribbon mode and the selected C_α_ atoms are shown as spheres. The arrows indicate the direction of the rotation relative to the static scaffold domain. (**B**) Proposed movement of the transport domain during transport. The full trajectory consists of two hinge motions with the elevator motion in between. The elevator motion has been proposed previously (refs 31, 32), and the hinge motion of the transport domain in the OFC is described in **Figure S6**. The static scaffold domain is depicted as a blue square and the transport domain is drawn as orange or pink rectangular. The grey boxes indicate the membrane.

## Conclusion

Compelling evidence has shown that BbZIP is an elevator-type transporter, but the detailed conformational changes and how metal substrate flows through the transport pathway remain to be elucidated. By locking the highly dynamic transporter into a fixed conformational state via Hg^2+^-mediated cross-linking, we solved a high-resolution crystal structure of BbZIP in a new IFC. A conserved metal relay in the water-filled metal release pathway was revealed, where a high-affinity metal binding site (M_ex_) at the exit of the pathway was identified. Mutagenesis and transport assays indicated that the M_ex_ site in BbZIP and the putative equivalent in hZIP4 negatively regulated the transport rate, likely by acting as a “metal sink” to reduce the overall metal transport rate. We speculate that it represents a potential regulatory mechanism. The identification of D604 in hZIP4 as an essential metal chelating residue indicated the importance of the putative transient metal binding site along the metal relay, which appears to be a conserved feature in some human ZIPs. The upward rotation of the transport domain relative to the scaffold domain, which is enabled by the Hg^2+^-mediated cross-linking, not only provided a new evidence for the elevator-type transport mode but more importantly revealed a continuous hinge motion of the transport domain.

The proposed trajectory consisting of a sequential hinge-elevator-hinge movement distinguishes BbZIP from other established elevator transporters and warrants further structural and biochemical/biophysical characterization to better elucidate the unique transport mechanism.

## Materials and Methods

### Genes, plasmids, mutagenesis, and reagents

The DNA encoding BbZIP (National Center for Biotechnology Information reference code: WP_010926504) was synthesized with optimized codons for *Escherichia coli* (Integrated DNA Technologies) and inserted into the pLW01 vector with a thrombin cleavage site inserted between the N-terminal His-tag and BbZIP. The DNA encoding human ZIP4 (GenBank access number: BC062625) from Mammalian Gene Collection were purchased from GE Healthcare, and inserted into a modified pEGFP-N1 vector (Clontech) in which the downstream EGFP gene was deleted and an HA tag was added at the C-terminus. Site-directed mutagenesis was conducted using the QuikChange® site-directed mutagenesis kit (Agilent). 1-Oleoyl-rac-glycerol (monoolein), N-ethylmaleimide (NEM), 1,10-phenanthroline, Tris (2-carboxyethyl) phosphine (TCEP) were purchased from Sigma-Aldrich.

### Protein expression, purification, and cross-linking by HgCl_2_

Expression of the A95C/A214C variant of BbZIP was the same as reported (*30, 32*). In brief, the expression was conducted in the *E*.*coli* strain C41 (DE3) pLysS (Lucigen) in LBE-5052 autoinduction medium for 24 hours at room temperature. After harvest, spheroplasts were prepared and lysed in the buffer containing 20 mM Hepes (pH 7.3), 300 mM NaCl, 0.25 mM CdCl_2_, and the cOmplete protease inhibitors (Sigma-Aldrich). n-Dodecyl-β-D-maltoside (DDM, Anatrace) powder was added to solubilize the membrane fraction (final concentration 1.5%, w/v). The His-tagged protein was purified using HisPur Cobalt Resin (Thermo Fisher Scientific) in 20 mM Hepes (pH 7.3), 300 mM NaCl, 5% glycerol, 0.25 mM CdCl_2_, and 0.1% DDM. The sample was then concentrated and loaded onto a Superdex Increase 200 column (GE Healthcare) equilibrated with the gel filtration buffer containing 10 mM Hepes, pH 7.3, 300 mM NaCl, 5% glycerol, 0.25 mM CdCl_2_, and 0.05% DDM. 1 mM TCEP was included throughout the purification process except for the size exclusion chromatography. To reduce Cd^2+^ level, the pooled peak fractions were briefly treated with 5 mM EDTA and immediately applied to a PD-10 desalting column (Cytiva) which has been equilibrated with the gel filtration buffer with Cd^2+^ excluded. The protein was then mixed with HgCl_2_ at a 1:10 molar ratio and allowed to incubate at room temperature for 30 minutes before setting up crystallization trays.

### Crystallization, data collection, and structure determination

Purified A95C/A214C variant was concentrated to 15 mg/ml and then mixed with the molten monoolein with two coupled syringes at a ratio of 2:3 (protein/monoolein, v/v). All crystallization trials were set up using a Gryphon crystallization robot (Art Robbins Instruments). 50 nl of BbZIP-monoolein mixture covered with 800 nl of well solution was sandwiched with lipidic cubic phase sandwich set (Hampton Research). Stick-shaped crystals appeared after one week under the condition containing 30% PEG200, 100 mM Hepes, pH 7.0, 100 mM NaCl, 100 mM CaCl_2_, at 21°C and grew to full size in two weeks. Crystals were harvested with a MiTeGen micromesh and flash-frozen in liquid nitrogen.

The X-ray diffraction data were collected at the General Medicine and Cancer Institutes Collaborative Access Team (GM/CA-CAT) (23-ID-B/D) at Advanced Photon Source (APS). The diffraction datasets were indexed, integrated, and scaled in HKL2000 v712. The apo state structure was solved in molecular replacement using the previously solved structure (PDB: 5TSB) as the search model in Phenix v1.10.1. Iterative model building and refinement were conducted in COOT v0.8.2 and Phenix, respectively. All figures of protein structures were generated by PyMOL v1.3 (Schrödinger LLC).

### Repeat-swap homology modeling to generate the OFC model

The modified protocol for repeat-swap homology model was used to generate the OFC models (*46*). In brief, the amino acids of the aligned TMs (α1-α3 vs. α6-α8, and α4 vs. α5) were swapped between the corresponding elements to generate a model sequence, which was then aligned with the wild type BbZIP (Figure S6A). To generate homology models, the sequence of the model was submitted to the SWISS-MODEL website (https://swissmodel.expasy.org/interactive#structure) and the structures of 8CZJ (metal-free IFC) and the new IFC structure reported in this work were used as template. In the energy minimized homology models, the loops and other regions not aligned between the model and BbZIP were trimmed and the amino acid registers were manually restored in COOT to obtain the final OFC models (Figure S6B, C).

### Growth-based functional assessment of BbZIP and Western blot

*E*.*coli* cells (C43 strain) transformed with the empty vector or the plasmids harboring the genes encoding BbZIP or the variants were allowed to grow in the autoinduction media overnight before they were applied on LB agar plates containing indicated concentrations of ZnCl_2_. After 24 hours of incubation at 37°C, the plates were imaged. To estimate expression of BbZIP and the variants, cell lysates were mixed with SDS sample loading buffer, applied to SDS-PAGE, and transferred to a PVDF membrane. After being blocked with 5% nonfat dry milk, the membrane was incubated with a mouse custom anti-BbZIP monoclonal antibody at 0.06 μg/ml at 4°C overnight (*32*). The bound primary antibodies were detected with an HRP-conjugated anti-mouse immunoglobulin-G at 1:5000 dilution (Cell Signaling Technology, Catalog# 7076S) by chemiluminescence (VWR). The images of the blots and plates were taken using a Bio-Rad ChemiDoc Imaging System.

### Mammalian cell culture, transfection, Cell-based zinc transport assay, and Western blot

Human embryonic kidney cells (HEK293T, ATCC, Catalog# CRL-3216) were cultured in Dulbecco’s modified eagle medium (DMEM, Thermo Fisher Scientific) supplemented with 10% (v/v) fetal bovine serum (FBS, Thermo Fisher Scientific) and Antibiotic-Antimycotic solution (Thermo Fisher Scientific) at 5% CO_2_ and 37°C. Cells were seeded on the polystyrene 24-well trays (Alkali Scientific) for 16 h in the basal medium and transfected with 0.8 μg DNA/well using lipofectamine 2000 (Thermo Fisher Scientific) in DMEM with 10% FBS.

The zinc transport activities of ZIP4 and the variants were tested using the cell-based transport assay. Twenty hours post transfection, cells were washed with the washing buffer (10 mM HEPES, 142 mM NaCl, 5 mM KCl, 10 mM glucose, pH 7.3) followed by incubation with Chelex-treated DMEM media (10% FBS). 5 μM Zn^2+^ (0.05 μCi/well) was added to cells. After incubation at 37°C for 30 min, the plates were transferred on ice and the ice-cold washing buffer with 1 mM EDTA was added to stop metal uptake. The cells were washed twice and pelleted through centrifugation at 120 x g for 5 min before lysis with 0.5% Triton X-100. A Packard Cobra Auto-Gamma counter was used to measure radioactivity. The transport activity was determined by subtracting the radioactivities of ^65^Zn associated with the cells transfected with the empty vector from those associated with the cells transfected with metal transporters.

hZIP4-HA and the variants expressed on the plasma membrane were determined by cell surface bound anti-HA antibody as reported (*39*). In brief, cells were washed twice with Dulbecco’s phosphate-buffered saline (DPBS) on ice and then fixed with 4% Formaldehyde for 5 min. Cells were then washed three times in DPBS and incubated with 5 μg/ml anti-HA antibody (Invitrogen, Catalog# 26183) at 4°C overnight. Cells were washed five times in DPBS to remove unbound antibodies, lysed in SDS-PAGE sample loading buffer, and eventually applied to SDS-PAGE and Western blot.

For Western blot, the samples mixed with the SDS sample loading buffer were heated at 96°C for 10 min before loading on SDS-PAGE gel. The protein bands were transferred to PVDF membranes (Millipore). After being blocked with 5% nonfat dry milk, the membranes were incubated with a mouse anti-HA antibody at 1:5000 dilution (Invitrogen, Catalog# 26183) at 4°C overnight. As loading control, β-actin levels were detected using a rabbit anti-β-actin antibody at 1:5000 dilution (Cell Signaling Technology, Catalog# 4970S). Primary antibodies were detected with an HRP-conjugated anti-mouse immunoglobulin-G at 1:5000 dilution (Cell Signaling Technology, Catalog# 7076S) for ZIP4 or an HRP-conjugated anti-rabbit immunoglobulin-G for β-actin at 1:5000 dilution (Cell Signaling Technology, Catalog# 7074S) by chemiluminescence (VWR). The images of the blots were taken using a Bio-Rad ChemiDoc Imaging System.

## Acknowledgments

We thank the beamline scientists at GM/CA-CAT at APS for the assistance in data collections. We thank the following funding that supported this work: NIH GM129004 and GM140931 (to J.H.).

## Author Contributions

J.H. conceived and designed the project. T.Z., Y.Z., and D.S. conducted experiments. T.Z., Y.Z., and J.H. analyzed the data and wrote the manuscript.

## Data and Material Availability

The atomic coordinates and structure factors have been deposited in the Protein Data Bank with the access code of 8J1M (https://www.rcsb.org/structure/unreleased/8J1M). All data needed to evaluate the conclusions in the paper are present in the main text and/or in the Supplementary Information. Additional data related to this paper may be requested from the authors.

The custom anti-BbZIP monoclonal antibody is available for research purpose upon request.

