## Supplementary material for "High-resolution structure of a mercury cross-linked ZIP metal transporter reveals delicate motions and metal relay for regulated zinc transport": SI

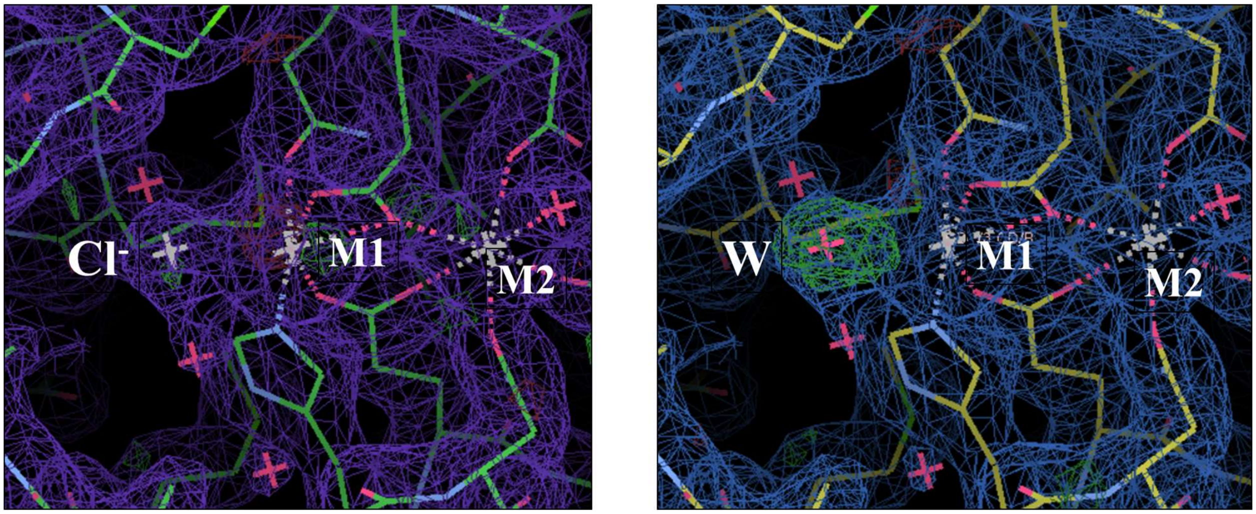


**Figure S1.** Electron density of the chloride ion coordinating $Cd^{2+}$ bound at the M1 site. *Left*: the density map with a modeled chloride ion ($Cl^-$). *Right*: density map with a modeled water molecule (W). Blue/purple meshes: 2Fo-Fc map at σ=1; Green meshes: positive densities in the Fo-Fc map at σ=3.

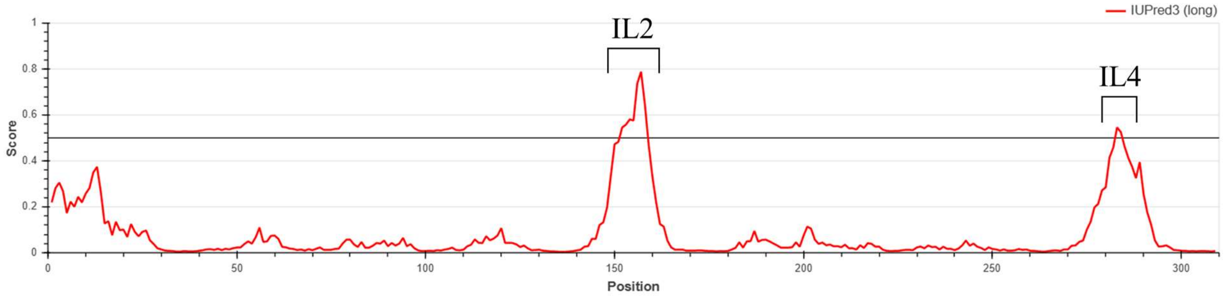


**Figure S2.** Prediction of intrinsically disordered region in BbZIP by IUPred3. The segment corresponding to the IL2 is predicted to be intrinsically disordered. The segment connecting TM7 and TM8 (IL4) is also predicted to be intrinsically disordered but to a less extent.

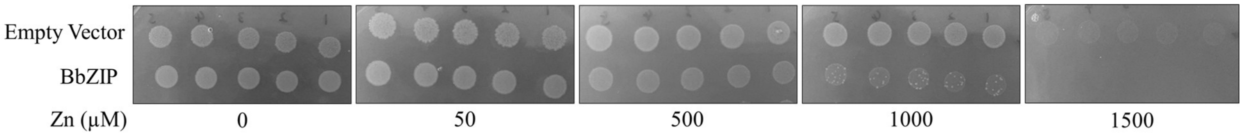


**Figure S3.** Growth of *E.coli* cells (C43 strain) with or without expressing BbZIP on LB agar plates containing indicated concentrations of $ZnCl_2$. The cells were allowed to grow on the agar plates at 37°C for one day. Expression of BbZIP suppresses cell growth only at high zinc concentrations (greater than 500 μM).

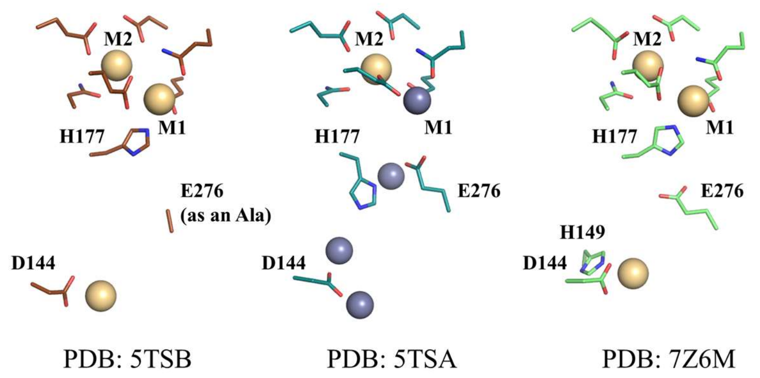


**Figure S4.** D144 and E276 in the metal release pathway in the previously solved Cd-bound structures. $Cd^{2+}$ and $Zn^{2+}$ ions are shown as light-brown and grey spheres, respectively.

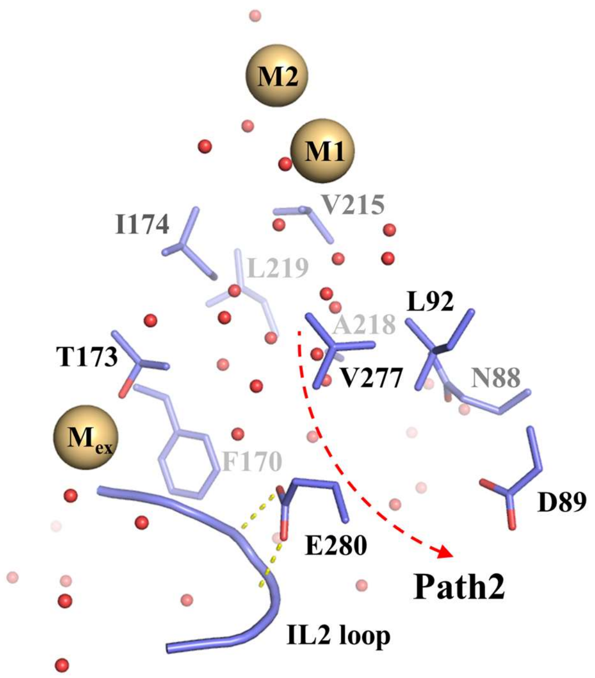


**Figure S5.** Residues along Path2 (the alternative metal release pathway). Most of the residues at the beginning of Path2 (as indicated by the red dashed arrow) are hydrophobic residues, likely resulting in a high energy barrier that prevents metal from entering. E280 forms two hydrogen bonds (yellow dashed lines) with the IL2 loop, which is as such not fully available for metal binding.

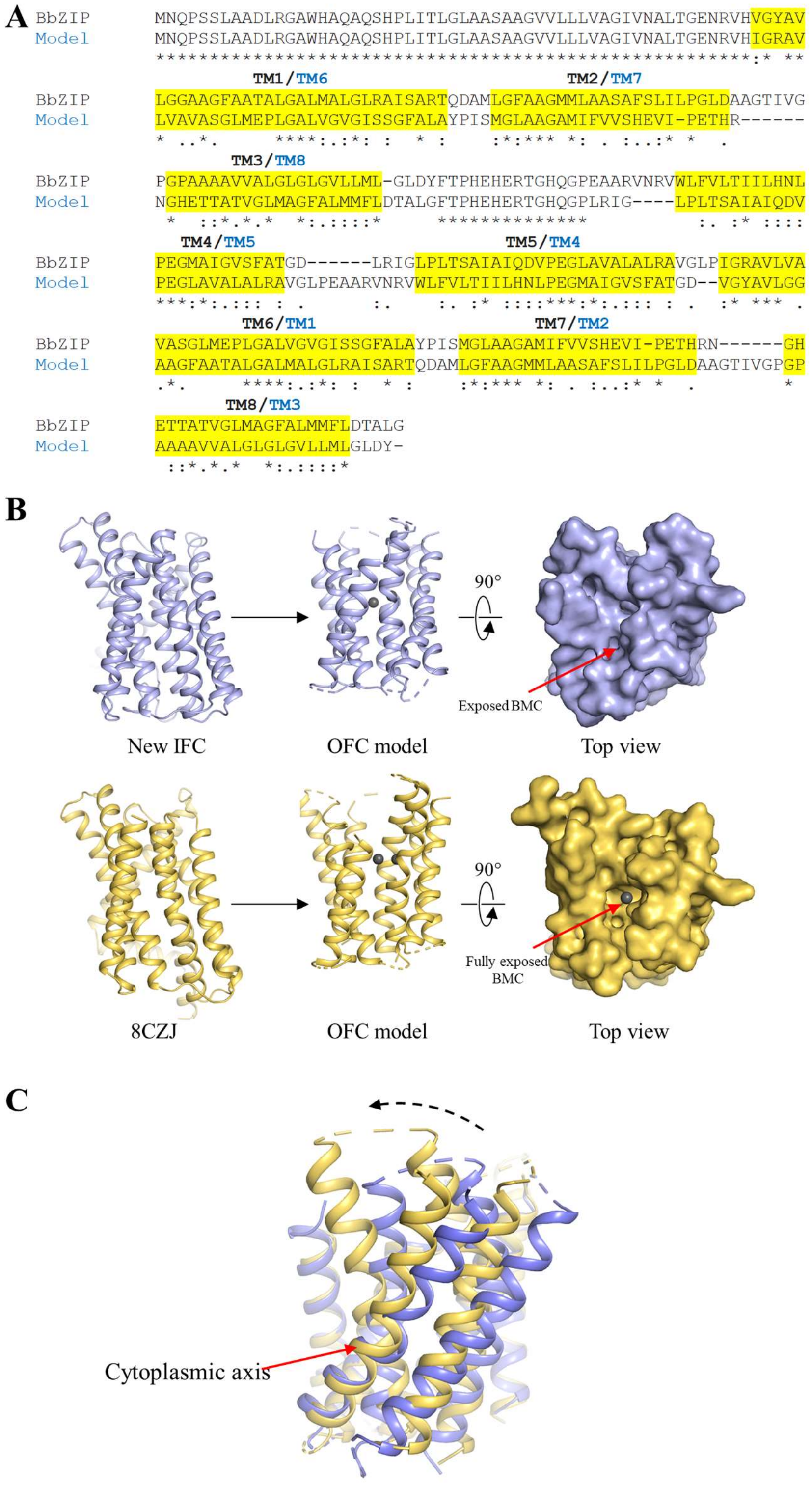


**Figure S6.** Generation and comparison of the OFC models. (**A**) Sequence alignment for generation of the OFC models using repeat-swap homology modeling. The symmetrically related elements are labeled on top of the sequences. (**B**) Templates (*left*) and the generated OFC models (*middle*). The top views in surface mode (*right*) show the entrance of the transport pathway. The BMC in the OFC model derived from 8CZJ (metal-free state) is more exposed than that from the new structure reported in this work. Metals bound at the BMC are depicted as grey spheres. (**C**) Structural comparison of the two OFC models. A hinge motion of the transport domain around the cytoplasmic axis (indicated by the dashed arrow) is revealed when the scaffold domains are structurally aligned.

**Table S1.** Crystallographic statistics

| **Crystal** | $Hg^{2+}$-crosslinked A95C/A214C BbZIP |
|---|---|
| **Data collection** | |
| Beamline | GM/CA-CAT (23-ID-B) |
| Wavelength (Å) | 0.987 |
| Space group | C $2_1$ |
| Unit cell | |
| a, b, c (Å) | 116.4, 51.0, 51.8 |
| α, β, γ (°) | 90, 113.9, 90 |
| [a]Resolution (Å) | 30.4 - 1.95 (2.02 – 1.95) |
| [a]Redundancy | 10.3 (2.3) |
| [a]Completeness (%) | 97.0 (78.4) |
| [a]$I/\sigma I$ | 14.3 (0.9) |
| [a,b]$R_{merge}$ | 0.172 (0.769) |
| [a,c]$R_{pim}$ | 0.05 (0.475) |
| [d]$CC_{1/2}$ of the highest resolution shell | 0.566 |
| **Refinement** | |
| Unique reflections | 19668 |
| Number of Atoms | 2217 |
| Protein | 2028 |
| Ligands | 131 |
| $H_2O$ | 58 |
| [e]$R_{work}/R_{free}$ | 0.1712/0.1999 |
| *Wilson B-factor* (Å$^2$) | 40.71 |
| *B*-factors (Å$^2$) | |
| Protein | 39.32 |
| Ligands | 59.94 |
| $H_2O$ | 45.9 |
| R.m.s. deviations | |
| Bond lengths (Å) | 0.007 |
| Bond angles (°) | 0.94 |
| Ramachandran plot (%) | |
| Favored | 97.91 |
| Allowed | 2.06 |
| Outliers | 0.0 |

[a]Highest resolution shell is shown in parentheses.

[b]$R_{merge} = \sum_{hkl} \sum_j |I_j(hkl)-\langle I(hkl)\rangle| / \sum_{hkl} \sum_j I_j(hkl)$, where $I$ is the intensity of reflection.

[c]$R_{pim}=\sum_{hkl} [1/(N-1)]^{1/2} \sum_j |I_j(hkl)-\langle I(hkl)\rangle| / \sum_{hkl} \sum_j I_j(hkl)$, where N is the redundancy of the dataset.

[d]$CC_{1/2}$ is the correlation coefficient of the half datasets.

[e]$R_{work} = \sum_{hkl} | |F_{obs}| - |F_{calc}| | / \sum_{hkl} |F_{obs}|$, where $F_{obs}$ and $F_{calc}$ is the observed and the calculated structure factor, respectively. $R_{free}$ is the cross-validation R factor for the test set of reflections (5% of the total) omitted in model refinement.
